# Resilience to cognitive aging is associated with responsiveness of dentate neurons generated throughout adult life

**DOI:** 10.1101/290676

**Authors:** Marie-Françoise Montaron, Vanessa Charrier, Nicolas Blin, Pierre Garcia, Djoher Nora Abrous

## Abstract

During aging some individuals are resilient to the decline of cognitive functions whereas others are vulnerable. These inter-individual differences in memory abilities have been associated with differences in the rate of hippocampal neurogenesis measured at old age. Whether the maintenance of the functionality of neurons generated throughout adult life is linked to resilience to cognitive aging remains completely unexplored. Using the immediate early gene Zif268, we analysed the activation of dentate granule neurons born in adult (3 month-old), middle-aged (12 month-old) or senescent (18 month-old) rats (n=96) in response to learning when animals reached 21 month-old. The activation of neurons born during the developmental period was also examined. We show that neurons generated 4, 10 or 19 months *before* learning (and not developmentally born neurons) are activated in senescent rats with good learning abilities. In contrast, aged rats with bad learning abilities do not exhibit an activity-dependent regulation of Zif268. In conclusion, we propose that resilience to cognitive aging is associated to the responsiveness of neurons born during adult-life. These data add to our current knowledge by showing that the aging of memory abilities stems not only from the number but also from the responsiveness of adult-born neurons.

## INTRODUCTION

Brain and cognition change with age, and although patterns of decline are evident at the population level, the rates of change differ among individuals as well as across brain regions and cognitive domains (Gray and Barnes, 2015; Nyberg *et al.*, 2012). Indeed, some old individuals exhibit cognitive abilities similar to those of younger ones (optimal/successful aging) whereas others show a clear substantial (suboptimal/accelerated aging) cognitive decline without signs of pathologies. Episodic memory is particularly sensitive to aging and investigations conducted so far have revealed both in humans and in animal models, that the preservation of episodic memory abilities is correlated to the structural and functional integrity of the hippocampal formation (Bettio *et al.*, 2017; Gonzalez-Escamilla *et al.*, 2018; Samson and Barnes, 2013). Several models and theories (maintenance, reserve, compensation) emerged in an effort to account for variability in cognitive outcome across old subjects and high level of neural plasticity has been proposed for brain reserve and resilience to cognitive aging (Nyberg *et al.*, 2012).

The ability of the adult brain, and in particular the dentate gyrus (DG) of the hippocampus, to create new neurons is a peculiar form of plasticity to protect the aging brain. Briefly, new dentate granule neurons (DGNs) generated throughout the entire life of an individual (Altman, 1962; Gross, 2000), humans included (Eriksson *et al.*, 1998; Moreno-Jiménez *et al.*, 2019; Spalding *et al.*, 2013), are integrated into functional circuits and play a crucial role in complex forms of learning and memory (Clelland *et al.*, 2009; Dupret *et al.*, 2008; Sahay *et al.*, 2011; Tronel *et al.*, 2012). In addition, both the addition and the elimination of new neurons in young adult rodent *before, during* or *after* learning are important for learning, remembering and forgetting (Akers *et al.*, 2014; Dupret *et al.*, 2007; Trouche *et al.*, 2009).

During aging, the rate of cell proliferation (and thus neurogenesis) decreases (Abrous *et al.*, 2005), a process associated to the progressive loss of neural stem cells (NSCs), to phenotypic and functional change of NSCs (Martin-Suarez *et al.*, 2019; Schouten *et al.*, 2019), or their niche (Diaz-Moreno *et al.*, 2018). Inter-individual differences in the rate of adult neurogenesis (ANg) has been linked to variability in spatial memory abilities of senescent animals: preserved memory functions are associated with the maintenance of a relatively high neurogenesis level measured *after* learning whereas memory deficits are linked to exhaustion of neurogenesis (Drapeau *et al.*, 2003). Moreover, we have found that corticosterone dampening from middle age has a beneficial effect on the rate of neurogenesis and spatial memory measured once animals have reached senescence (Montaron *et al.*, 2006). Together, this last set of data raises the fascinating hypothesis that neurons generated throughout adult life could constitute a mechanism that promotes resilience to cognitive aging.

To tackle this question, we examined the activation of DGNs generated throughout adult life in the maintenance of memory function by imaging them when animals reached senescence. DGNs born in adult (3 month-old), middle-aged (12 month-old) or senescent (18 month-old) rats were labeled with analogs of thymidine and their activation in response to spatial learning was measured using Zif268, an Immediate Early Gene (IEG) (Tronel *et al.*, 2015b), when animals have reached senescence. The activation of DGNs born during development was also examined.

## MATERIALS AND METHODS

### Animals

For these experiments, a total of 96 male Sprague-Dawley rats (OFA, Janvier, France) were used. Animals were housed collectively until behavioural testing under a 12h:12h light/dark cycle with *ad libitum* access to food and water. Temperature (22°C) and humidity (60%) were kept constant.

*In the first experiment*, male rats (n=19), were 16-month-old on delivery. *In the second experiment*, rats (n=32) were 2-month-old on delivery. *In the third and fourth experiments*, rats (n=25) were 21 day-old on delivery. In the fifth experiment, pregnant Sprague-Dawley female rats (n=4) were individually housed in transparent cages. After delivery, the litters were raised by their biological mothers until weaning (21 days after birth). After weaning, only the male progeny (n=20) was kept. Rats were individually housed before the beginning of behavioral training. Animals with a bad general health status or tumors were excluded. Experimental procedures have been planned respecting the European directive of the parliament and the conceal of September 22, 2010 (2010/63/UE, 5012006A).

### Thymidine analogue injections

Newly-born cells were labeled by the incorporation of synthetic thymidine analogues (XdU, Sigma Aldrich, Saint Louis, USA Table 1). *In the first experiment*, rats were injected with 5-bromo-2’-deoxyuridine (BrdU) according to a previously described protocol (Drapeau *et al.*, 2003; Drapeau *et al.*, 2007). These animals received one daily BrdU injection (50 mg/kg/day; ip) for five days when 18-month-old, i.e. four months before training. *In the second experiment*, rats received five injections of 5-chloro-2’-deoxyuridine (CldU) when 3-month-old and five injections of 5-iodo-2’-deoxyuridine (IdU) when 12-month-old(Dupret *et al.*, 2007), both at equimolar doses of 50mg BrdU/kg. *In the third and fourth experiments*, animals received one injection of CldU when 28 day-old (equimolar dose of 50mg BrdU/kg). *In the fifth experiment*, pregnant female rats received two injections of 5-chloro-2’-deoxyuridine (CldU, equimolar dose of 50mg BrdU/kg 50mg/kg) at E18.5 and E19.5.

**Table 1:** Summary of the procedures

| Batch | Experiment | XdU | Rats' age at XdU injections | Rats' age at time of sacrifice | Neurons' age at time of sacrifice | Group size |
| --- | --- | --- | --- | --- | --- | --- |
| 1 | Recruitment of Adu-DGNs of 4-month-old | BrdU | 18 months | 22 months | 4 months | C = 5<br>AI = 7<br>AU=7 |
| 2 | Recruitment of 10-month-old Adu-DGNs | IdU | 12 months | 22 months | 10 months | C = 10<br>AI = 11<br>AU=11 |
|  | Recruitment of 19-month-old Adu-DGNs | CldU | 3 months | 22 months | 19 months | C = 10<br>AI = 11<br>AU=11 |
| 3 | Recruitment of DGNs born in adolescent rats | CldU | PN28 | 22 months | 21 months | AI = 5<br>AU=5 |
| 4 | Recruitment of DGNs born in adolescent rats | CldU | PN28 | 14 months | 15 months | AI = 5<br>AU=5 |
| 5 | Recruitment of DGNs born in embryos | CldU | Ed18.5 | 15 months | 15 months | AI = 6<br>AU=7 |

### Water-maze training

Rats were tested in the water-maze when 22-month-old (experiments 1,2,4) or 15-month-old (experiments 3,5). The apparatus consisted of a circular plastic swimming pool (180 cm diameter, 60 cm height) that was filled with water (20 ± 1°C) rendered opaque by the addition of a white cosmetic adjuvant. Before the start of training, animals were habituated to the pool for two days for one minute per day. During training, the *Learning* group (L) was composed of animals that were required to locate the submerged platform, which was hidden 1.5 cm under the surface of the water in a fixed location, using the spatial cues available within the room. Rats were all trained for four trials per day (90 s with an inter-trial interval of 30 s and released from 3 different starting points that varied randomly each day). If an animal failed to locate the platform, it was placed on that platform at the end of the trial. The time to reach the platform was recorded using a video camera that was secured to the ceiling of the room and connected to a computerised tracking system (Videotrack, Viewpoint). Daily results were analyzed in order to rank animals according to their behavioral score calculated over the last 3 days of training (when performances reached an asymptotic level). The behavioral scores calculated over the whole training duration of Aged unimpaired (AU) rats were below the median of the group whereas those of Aged Impaired (AI) animals were above the median of the group. Control groups consisted of animals that were transferred to the testing room at the same time and with the same procedures as trained animals but that were not exposed to the water maze.

### Immunohistochemistry

Animals were sacrificed 90 min after the last trial (Table 1). The different age-matched control groups were sacrificed within the same period. Free-floating sections (50 µm) were processed using a standard immunohistochemical procedure to visualize the thymidine analogs (BrdU, CldU, IdU) on alternate one-in-ten sections using different anti-BrdU antibodies from different vendors (for BrdU: 1/200, Dako, Glostrup, Denmark; CldU: 1/500, Accurate Chemical, Westbury, USA; IdU: 1/200, BD Biosciences, San Jose, USA) and Zif268 (1:500, Santa Cruz Biotechnology, Dallas, USA). The number of XdU-immunoreactive (IR) cells in the granule and subgranular layers (gcl) of the DG was estimated on a systematic random sampling of every tenth section along the septo-temporal axis of the hippocampal formation using a modified version of the optical fractionator method. Indeed, all of the XdU-IR cells were counted on each thick section and the resulting numbers were tallied and multiplied by the inverse of the section sampling fraction (1/ssf=10 for BrdU and IdU-cells that were counted in both sides of the DG, 1/ssf=20 for CldU-IR cells that were counted in the left side). The number of Zif268-IR cells (left side) was determined using a 100x lens, and a 60 µm x 60 µm frame at evenly spaced x-y intervals of 350 µm by 350 µm with a Stereo Investigator software (Microbrightfield).

### Activation of new cells

The activation of adult-born cells was examined using immunohistofluorescence. To visualize cells that incorporated thymidine analogues, one-in-ten sections were incubated with different anti-BrdU antibodies (BrdU & CldU, rat primary antibodies at 1/200 Accurate Chemical; IdU, mouse primary antibodies at 1/200, BD Biosciences). Sections were also incubated with Zif268 (rabbit,1:500, Santa Cruz Biotechnology). Bound antibodies were visualized respectively with Cy3-goat anti-rat (1:1000, Jackson, West Grove, USA) or Cy3-goat anti-mouse (1:1000, Jackson) and Alexa-488-goat anti-rabbit antibodies (1:1000, Jackson). CldU-Zif268 and IdU-Zif268 labeling were analyzed on different sections because of some cross reactivity between secondary antibodies made in mice or rat (Fig 1 in (Tronel *et al.*, 2015b)). All BrdU-, CldU- or IdU-labeled cells expressing Zif268 (one side) were determined using a confocal microscope with HeNe and Arg lasers (Leica, DMR TCSSP2AOBS), with a plane apochromatic 63X oil lens (numerical aperture 1.4; Leica). The percentage of BrdU-, CldU- or IdU-labelled cells that expressed Zif268 was calculated as follow: (Nb of XdU^+^/IEG^+^ cells)/[(Nb of XdU^+^/IEG^−^ cells) + (Nb of XdU^+^/IEG^+^ cells)] x 100. All sections were optically sliced in the Z plane using 1 µm interval and cells were rotated in orthogonal planes to verify double labelling.

**Figure 1.**
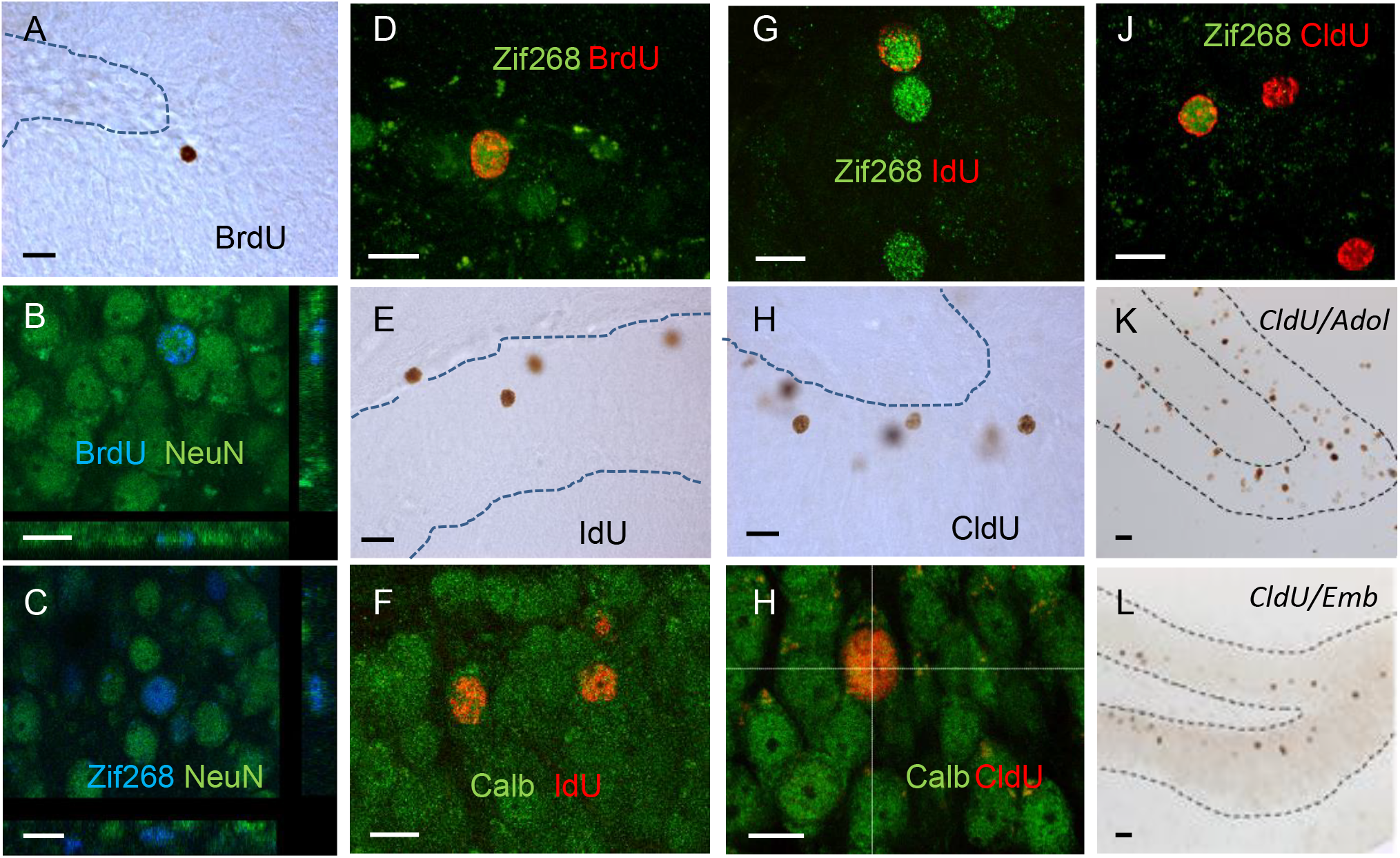
Granule neurons in the of aged DG. **(A)** Illustration of 4-month-old BrdU-IR neurons in an animal with preserved memory. **(B)** Confocal photomicrographs of 4-month-old BrdU-IR cells (blue) expressing NeuN (green). Confocal photomicrographs of **(C)** neurons (NeuN, green) expressing Zif268 (blue) and of **(D)** 4-month-old BrdU-IR cells (red) expressing Zif268 (green). **(E)** Illustration of 10-month-old IdU-IR neurons. Confocal photomicrographs of IdU-IR cells (red) expressing **(F)** Calbindin (green) or (**G**) Zif268 (green). Illustration of 19-month-old CldU-IR neurons. **(H)** Confocal photomicrographs of CldU-IR cells (red) expressing **(I)** Calbindin (green) or **(J)** Zif268 (green). **(K)** Illustration of CldU-IR neurons born in adolescent rats (PN28). **(L)** Illustration of CldU-IR neurons born in embryons (ED18.5). Bar scale for DAB= 20µm. Bar scale for confocal illustration =10µm.

### Analysis of phenotype

One-of-ten series was incubated with a rat monoclonal anti-BrdU antibody (1/200, Accurate Chemical) and with a mouse monoclonal anti-NeuN antibody (1:500, Millipore, Massachusetts, USA). Bound anti-BrdU and anti-NeuN antibodies were visualized with a Cy3-goat anti-rat (1:1000, Jackson) and an Alexa 488-goat anti-mouse IgG antibody (1:1000, Jackson). The phenotype of IdU-IR cells and CldU-IR cells was determined using rabbit anti-calbindin antibodies (1/200, Millipore) that were revealed with Alexa 488-goat anti-rabbit IgG antibodies (1/500, Jackson). We also analysed the phenotype of Zif268 cells by incubating one-in-ten sections with a rabbit anti-Zif268 antibody (1:500, Santa Cruz Biotechnology) and a mouse monoclonal anti-NeuN antibody (1:500, Millipore). Bound anti-Zif268 and anti-NeuN antibodies were visualized with a Cy3-goat anti-rabbit (1:1000, Jackson) and an Alexa 488-goat anti-mouse IgG antibody (1:1000, Jackson).

### Statistical analysis

Data (mean±s.e.m.) were analysed using an ANOVA or Student’s t-test (2 tails) when necessary.

## RESULTS

In a first step we sought out to determine whether new neurons born during ***senescence*** are recruited by spatial learning. To do so, eighteen-month-old rats were injected with BrdU according to a previously described protocol (Table 1) and were trained four months later in the water maze using a reference memory protocol (Drapeau *et al.*, 2003). Animals were trained for eleven days (**Figure S1A,B**) until the aged-unimpaired rats (AU) learned the task (day effect on the Latency: F_11,66_=2.35, p=0.016; day effect on Distance: F_11,66_=2.76, p=0.005) and reached asymptotic levels of performances (with no statistical significant differences between the last 3 days). In contrast, the aged-impaired (AI) rats did not learn the task although they were searching and finding the platform most of the time (2 or 3 trials out of 4) (day effect on the Latency: F_11,66_=1.25, p=1.25; day effect on Distance: F_11,66_=0.96, p=0.48). Ninety minutes after the last trial, animals (and their age-matched control group) were sacrificed for immunohistochemistry. At the time of sacrifice, BrdU-IR cells were 4 months-old and the majority was located within the granule cell layer (GCL) (Figure 1A).

These cells were more numerous in the GCL of aged animals with good learning abilities (AU) compared to aged animals with memory deficits (AI) (Figure 2A, F_2,16_=7.64, p=0.05 with C=AI<AU at p<0.01). This finding is consistent with our previous study showing that the number of neurons generated one month *after* learning is higher in AU compared to AI (Drapeau *et al.*, 2003) senescent rats. More than fifty percent of BrdU-IR cells in the GCL expressed NeuN (Figure 1B) and neuronal differentiation was not different among groups (Figure 2B, F_2,16_=2.07, p=0.15).

**Figure 2.**
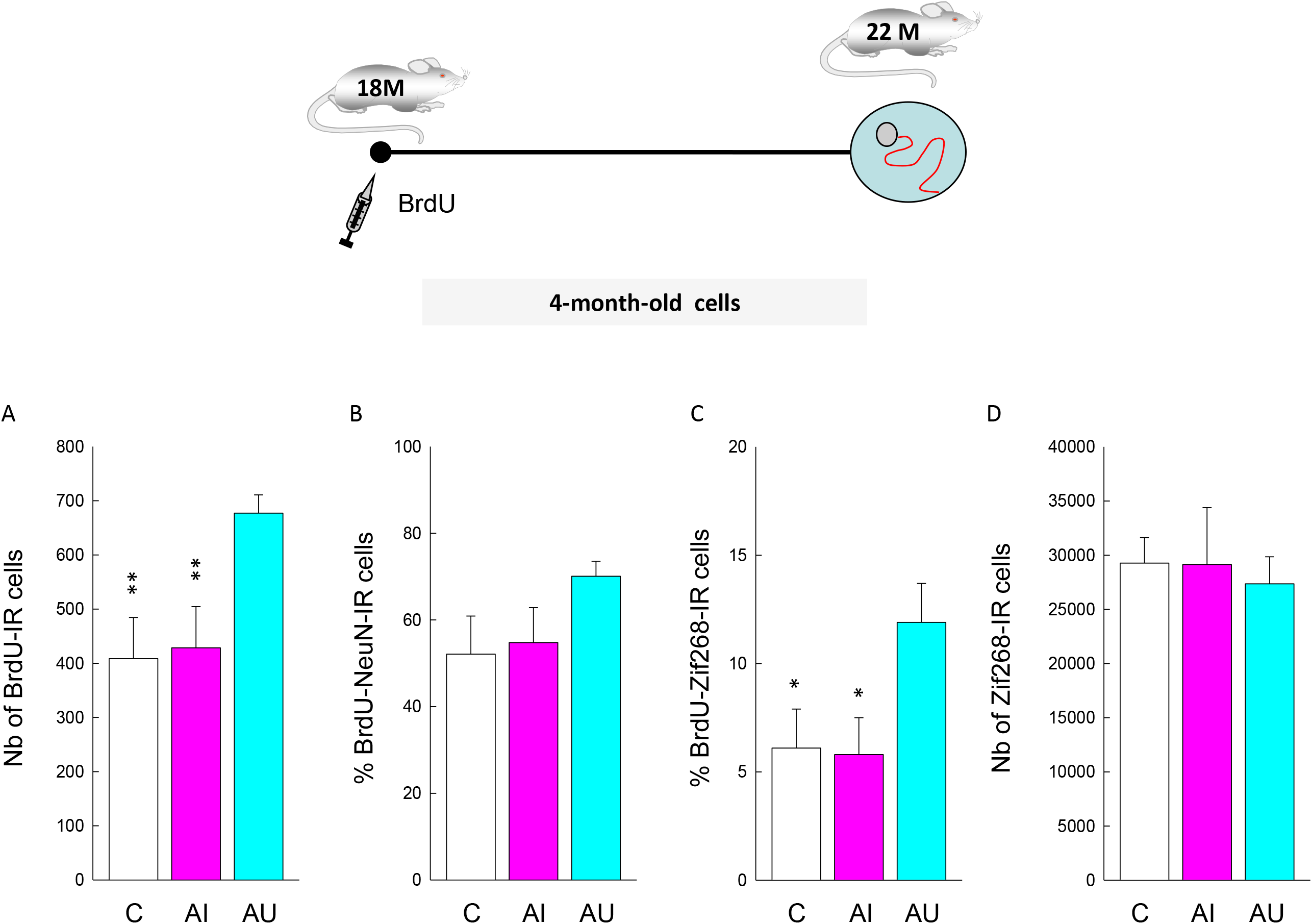
Neurons produced during old age are activated by spatial learning. Top: Experimental design. (**A**) The number of BrdU-IR cells is higher in the aged rats that learned the task (AU) compared to those with spatial memory deficits (AI) or to control animals (C). (**B**) The percentage of cells differentiating into neurons (BrdU-IR cells expressing NeuN) is similar between the three groups. (**C)** The expression of Zif268 in BrdU-IR cells generated in senescent DG is increased in AU compared to AI rats and C rats. (**D**) The number of neurons expressing Zif268 is similar between the three groups. *: p<0.05, **: p<0.01 compared to AU.

To determine whether newborn neurons are recruited by learning, we used Zif268 since this IEG is still expressed in the old DG (Gheidi *et al.*, 2013; Marrone *et al.*, 2011). Given that a substantial fraction of cells generated during senescence did not express NeuN, we verified in trained animals that Zif268 expressing cells were expressing NeuN (Figure 1C). We found that the vast majority of activated cells (Zif268) were neurons (NeuN) and that this ratio was similar between good and bad learners (AI: 96.4 ± 0.5; AU: 96 ± 1.3, p>0.05). Then we examined the activation of adult-born cells, meant to be neurons, in response to learning (Figure 1D). We found that the percentage of BrdU-IR cells expressing Zif268-IR in aged animals with good learning abilities was greater than that of aged animals with memory deficits and of untrained control groups (Figure 2C, F_2,16_=3.70, p=0.05 with C=AI<AU at p<0.05). In contrast, the total number of Zif268-IR nuclei did not differ between groups (Figure 2D, F_2,16_=0.25, p=0.78). These results show that neuronal cells in the senescent DG are recruited by spatial learning and not by nonspecific effects of training (swimming, stress) as revealed by the lowest level of recruitment of 4-month-old cells in aged impaired and control animals.

Then we asked whether neurons born earlier, i.e. in middle-age or young adulthood, are also recruited by learning during aging. For this purpose, animals were injected with CldU when 3-month-old, and with IdU when middle-aged (at twelve months old; Table 1). Animals were trained ten months later for eleven days until the AU learned the task (day effect on the Latency: F_10,100_=22.08, p<0.001; day effect on Distance: F_10,100_=18.77, p<0.001) and reached three days of stable performances (**Figure S1C,D)**. In this batch, the AI showed a dramatic improvement of their performances on the last training day (day effect on the Latency: F_10,100_=6.67, p<0.001; day effect on Distance: F_10,100_=22.08, p<0.001). Trained animals (and their age-matched control group) were sacrificed 90 minutes after the last trial. At the time of sacrifice IdU cells were 10-month-old (Figure 1D). Their number was not influenced by training or by the cognitive status of the animals (Figure 3A, F_2,29_=0.87, p=0.43). More than eighty percent of IdU cells expressed the neuronal marker calbindin (Figure 1F, 3B, F_2,28_=4.21, p=0.02 with C=AI<AU at p=0.02). The percentage of neurons born during middle-age and expressing Zif268 was greater in the AU group than that measured in AI and C groups (Figures 1G, 3C, F_2,29_=4.87, p=0.02 with C=AI<AU at p<0.01 and p<0.05 respectively).

**Figure 3.**
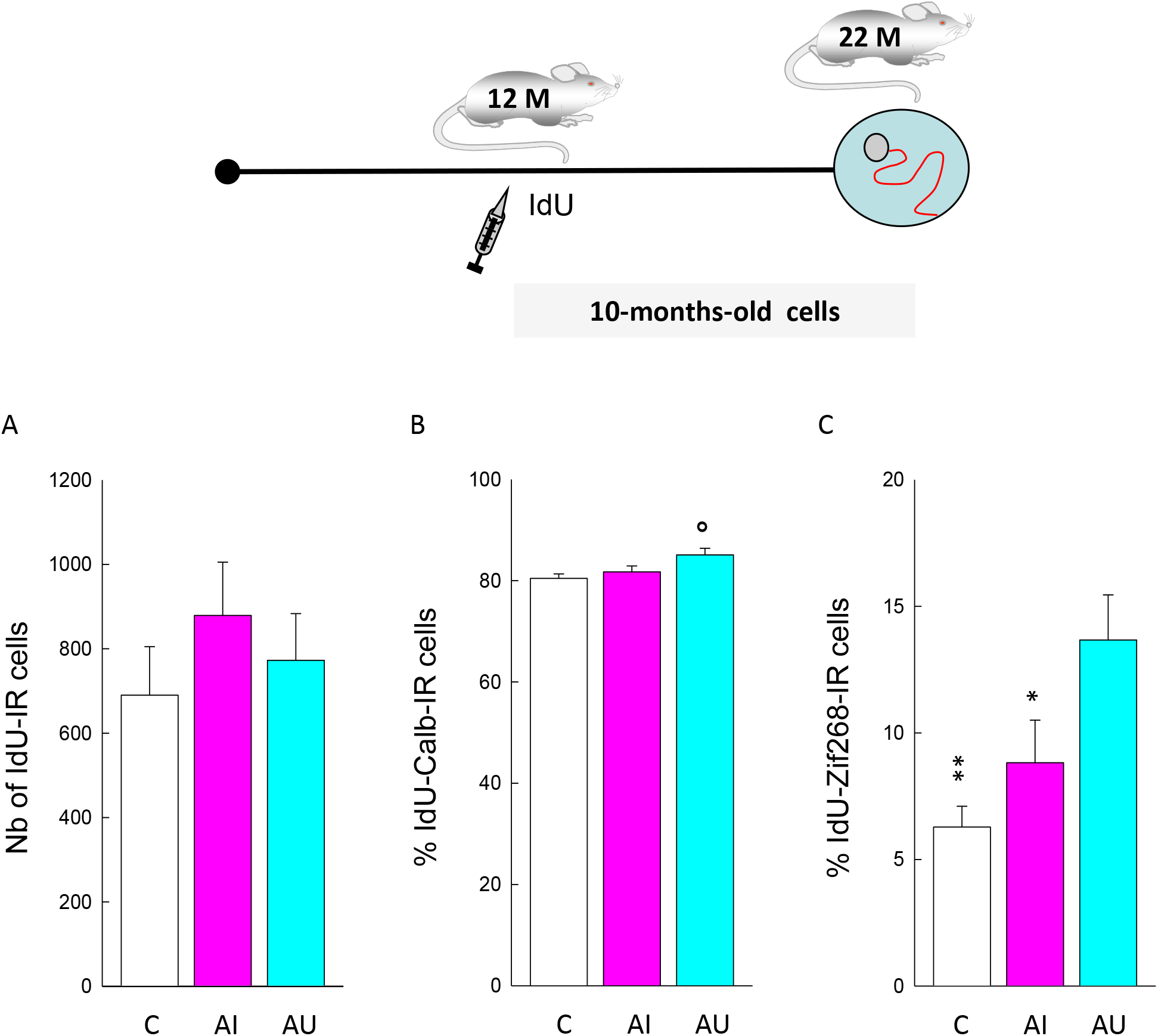
Neurons produced during middle-age are activated by spatial learning in aged good leaners. Top: Experimental design. (**A**) The number of IdU-IR cells -generated at mid-age-is independent of the memory abilities measured when rats reached senescence. (**B**) The percentage of cells differentiating into neurons (IdU-IR cells expressing Calbindin) is slightly increased in AI (compared to C and AU). (**C**) The expression of Zif268 in IdU-IR cells is increased in AU rats compared to AI rats and C rats.*: p<0.05, **: p<0.01 compared to AU. °: p<0.05 compared to C.

CldU-IR cells examined in the same animals were 19-month-old (Figure 1H). Their number was not influenced by training or the cognitive status of the animal (Figure 4A, F_2,29_=0.52, p=0.6). By analysing the phenotype of CldU cells with calbindin, we found that exposure to the water maze slightly increased neuronal differentiation (Figure 1J,4B, C: 82.8 ±.1% AI: 86.9 ± 0.8%; AU: 86.2 ± 0.7%; F_2,29_=6.54, p<0.01 with C<AI=AU at p=0.01). Again, we found that the percentage of CldU-IR cells expressing Zif268 was greater in the AU group than that measured in AI and C groups (Figures 1J, 4C, F_2,29_=6.96, p=0.004 with C=AI<AU at p<0.01 and p<0.05 respectively). The total number of cells expressing Zif268-IR (C: 29270.02 ± 2360.54: AI: 26068.94 ± 2366.78; AU: 28739.22 ± 3095.74, F_2,29_=0.42, p=0.65) did not differ between groups.

**Figure 4.**
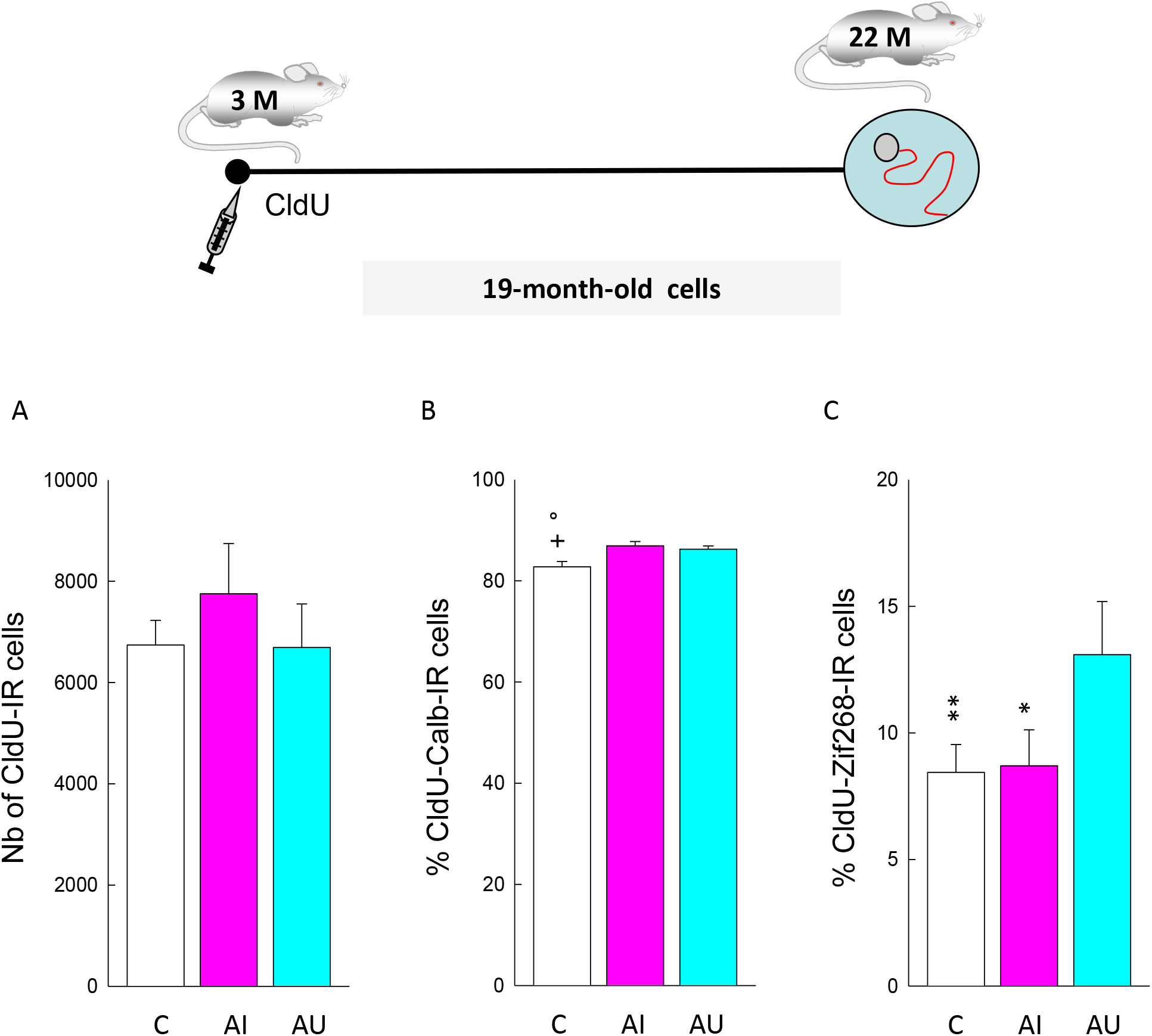
Neurons produced during young adulthood are activated by spatial learning in good leaners. Top: Experimental design. (**A**) The numbers of CldU-IR cells generated when animals are young adult is independent of the memory abilities measured when rats reached senescence. (**B**) The percent of CldU-IR cells expressing calbindin is increased by training. (**C**) The expression of Zif268 in CldU-IR cells generated in young adult DG is increased in AU compared to AI rats and C rats. °: p<0.05 compared to AU, +: p<0.05 compared to AI. *: p<0.05, **: p<0.01 compared to AU.

Finally, we explored the role of dentate granule born during development of the DG by tagging neurons born in adolescent rats (PN28, Figure 1K) and neurons born in embryos (E18.5, Figure 1L) with CldU. Animals were sacrificed when 22 or 15-month-old. In all conditions, the number of CldU-IR cells and the percentage of CldU-IR cells expressing Zif268 were similar in AU and AI (**Table S1**).

## DISCUSSION

To determine whether neurons generated during adult life participate to learning abilities in old age, the expression of the IEG Zif268 in new neurons was assessed. We found that cells generated during young adulthood, middle-age and senescence survive for a long period of time and are functionally integrated into the dentate network. When taking into account individual differences in memory abilities, we highlight that although the number of new cells generated in 12-month-old animals is decreased tenfold compared to 3-month-old rats, the total number of CdU-IR or IdU-IR cells measured when animals reached senescence is similar between AU and AI and not different from untrained control animals.

These conclusions have been obtained using two different cohorts of rats. The first one was utilized to study Adu-DGNs generated in senescent DG and the second one to study Adu-DGNs generated in young adult and middle aged DG. When comparing the behavior of the two batches of rats, it appears that deficits in the aged-impaired rats were much pronounced for the first batch of animals. This cohort effect, a well know phenomena in aging research (Schaie and Willis, 2015), could be related to the housing condition. Indeed, the first batch was raised in the vendor facilities until 16-month-of age whereas the second one was raised in-house. Supporting this, in our previous experiments performed in rats not aged in-house, the difference between AU and AI were more pronounced that observed in the second experiment (Desjardins *et al.*, 1997; Drapeau *et al.*, 2007; Schaie and Willis, 2015). However, independently of the cohort of rats or of adult-born neurons the same profile of activation was observed (C=AI <AU).

While the process of neurogenesis have been well characterized in young adult rodents (Aimone *et al.*, 2014), information about their aging and their function is less abundant (Drapeau and Abrous, 2008; Encinas and Fitzsimons, 2017; McAvoy and Sahay, 2017). The number of stem cells and the rate of cell proliferation dramatically decrease with age, along with their neuronal differentiation and the number of immature neurons (<4 weeks-old) is thus significantly decreased. Recently, the development and functional integration of these cells has been described to be delayed by age (Trinchero *et al.*, 2017). Indeed, 3-week-old neurons generated in middle-aged mice (10-14 months) displayed shorter and simpler dendrites and a dramatic reduction in spine number compared to 2-month-old mice, and exhibited immature neuronal electrophysiological properties as revealed by the lack of functional glutamatergic synaptic inputs (Trinchero *et al.*, 2017). In fact, their overall mature excitability and maximal glutamatergic connectivity is delayed compared to neurons born in younger animals as achieved within 10 weeks (Trinchero *et al.*, 2019). The long-term destiny of adult-born generated in young adult animals has not been explored in depth. We and others have shown that contrary to what was initially hypothesized (Gross, 2000), the new neurons survive for several months (Kempermann *et al.*, 2003; Tronel *et al.*, 2015b) and even years in the DG (present results) and do not show signs of decline in excitability when they age: 5-month-old neurons are as excitable as 1-month-old-cells; they can even exhibit high levels of excitability following either enriched environment exposure or induction of LTP (Ohline *et al.*, 2018). This latest very exciting result supports our hypothesis that even when several months-old, Adu-DGNs are still plastic, they do not retire and participate in memory functions (Abrous and Wojtowicz, 2015; Lemaire *et al.*, 2012; Tronel *et al.*, 2015b), and even more so their persistence is not passive, but a result of their activity.

Here, we found that between middle-age and senescence the number of cells is further decreased, but then a difference among the AU and AI groups appears. Based on previous data, it is likely that the emergence of such a difference results from a difference in cell proliferation (Diaz-Moreno *et al.*, 2018; Drapeau *et al.*, 2003), neuronal differentiation (Drapeau *et al.*, 2003; Qiao *et al.*, 2019), cellular senescence (Hernandez-Segura *et al.*, 2018; Micheli *et al.*, 2019), changes in the neurogenic niche (Fan *et al.*, 2017) and/or to the systemic milieu (Mahmoudi *et al.*, 2019; Smith *et al.*, 2015; Villeda *et al.*, 2011). Interestingly, we have shown that the senescent neurogenic niche is capable to rejuvenate upon removal of corticosterone (Montaron *et al.*, 1999) or addition of Sulfate pregnenologne (Mayo *et al.*, 2005) indicating that neural stem cells are not depleted and keep their abilities to divide.

The main finding of our study is that the ability for newborn cells to be recruited by learning in aged rats depends upon their memory abilities. Indeed, the percentage of adult-born cells expressing Zif268 was higher in animals that learned the task compared to animals that did it to a lesser extent. This finding is in accordance with our previous data showing that i) when compared to control rats (naïve rats or rats trained to find a visible platform), adults required to use an hippocampal-dependent strategy in the water-maze (or the dry maze) exhibit an increased percentage of mature adult-born neurons expressing Zif268 (Tronel *et al.*, 2015a; Tronel *et al.*, 2015b), and ii) ablating mature adult-born neurons generated four months before training (when animals where 3 months old) delays the ability of rats to learn such a task (Lemaire *et al.*, 2012). In the present experiment the percentage of adult-born cells expressing Zif268 in each experimental group was similar for the three neuronal populations studied. It was thus independent of the age of the animals at the time of labeling (3, 12, and18 months) and of the age of the cells at the time of training (4, 10, and 19 months). It was also independent of whether or not the total number of XdU cells differed between AU and AI groups. Note that even if in the first batch of animals, fifty percent of BrdU-IR cells differentiated into neurons, 96% of Zif268 cells were neurons suggesting that all BrdU-Zif268 cells were meant to be activated new neurons.

It could be argued that neurons born during development, which represent a major part of the DG, are also involved in differences in spatial memory abilities in old age. However, three arguments seem to rule out this hypothesis. First the total number of granule cells is similar between AU and AI groups (Drapeau *et al.*, 2003; Drapeau *et al.*, 2007; Rapp and Gallagher, 1996). Second, we have shown that neurons born in embryos, neonates, juveniles are not activated by spatial learning when they are mature compared to neurons of the same age born in adults. Indeed, the former are not recruited by spatial learning in the water maze when animals are tested at 7-month-old (Tronel *et al.*, 2015b). Third, if neurons generated during development (pre- and post-natal periods) were activated by spatial learning, given their high numbers, differences in the total number of Zif268 cells should have emerged as a function of the cognitive status. We began to explore their role in aging in the present manuscript and found that neurons born during the juvenile period (PN28) or the embryonic period (E18.5/19.5) are not differentially recruited in the good and bad learners.

One question that we did not address is whether the three neuronal populations studied participate to the same extent to learning. To address this point, sophisticated models that allow to selectively tag new neurons generated within a defined period of time (adulthood, middle-age or senescence) and to ablate them during training performed at senescence, are required. One possibility would be to take advantage of the recently developed pharmacogenetic approach of DREADD (Designer Receptor Exclusively Activated by Designer Drug)(Alvarez *et al.*, 2016) or optogenetic (Gu *et al.*, 2012) in order to tag specifically neurons born in young adult rats and manipulate them when animals have reached senescence.

A previous study has shown that 4-month-old neurons generated in old rats exhibiting spatial memory deficits are recruited in response to spatial exploration behavior with the same probability than 4-month-old neurons generated in aged good learners or in young adult rats (Marrone *et al.*, 2012). From this dataset it was concluded that disrupted information processing at old age may be linked to a reduced number of adult-generated granule cells, and not to a deficit in their functionality. However, in this study the activation of adult-generated neurons was evaluated in response to a simple form a learning (spatial exploration). Taking the present data into consideration, we rather suggest that adult-born neurons in AU are sufficiently connected to integrate simple stimulations generated during simple form of learning but insufficiently integrated to process the complex stimulations generated during spatial navigation.

Zif268 is known to be regulated in an activity-dependent manner by learning (for review see (Veyrac *et al.*, 2014)). It is overexpressed in response to different types of learning in distinct structures and circuits that are processing the ongoing information and several arguments indicate that it is required for the stabilization (and not acquisition) of long-lasting memories. Although the mechanisms are not fully understood, the activation of Zif268 may strengthen/stabilize the memory trace. It can be hypothesized that during learning the activation of Zif268 in adult-born neurons of GL may be involved in the formation, stabilization and reactivation of place cells in the hippocampal network, events known to support spatial learning (O’Keefe J, 1978).

Here we hypothesize that adult-born neurons that do not exhibited activity-dependent regulation of Zif268 become functionally silent in the course of aging, leading to memory deficits. Although the firing patterns that are sufficient to induce Zif268 in adult-born neurons in “behaving” animals are so far unknown, adult-born neurons silencing may have several origins. It may result from a loss of synaptic inputs (Fischer *et al.*, 1987; Geinisman *et al.*, 1986; Smith *et al.*, 2000) altering the ability to fire properly (Ahlenius *et al.*, 2009); these synaptic alterations of Adu-DGNs could be linked to the acceleration of senescence through epigenetic changes (Penner *et al.*, 2010; Penner *et al.*, 2011), decrease autophagy activity (Glatigny *et al.*, 2019) or changes of the local and systemic milieu (Fan *et al.*, 2017).

In conclusion, our results highlight the importance of neurons born throughout adult-life in providing resilience to age-related memory disorders. They reveal a novel perspective for developing therapies to promote resilience to age-related memory disorders or to rejuvenate the DG by acting throughout adult life on adult-born dentate neurons (Fan *et al.*, 2017; Mahmoudi *et al.*, 2019).

## ACKNOWLEDGEMENTS

The authors thank Dr M Koehl (Inserm, UMR 1215) and Dr A Schinder (Leloire Institute, BA, Argentina) for useful discussion. We greatly acknowledge C Dupuy for animal care and A Mangin for his technical help. This work was supported by Institut National de la Santé et de la Recherche Médicale, Centre National de la Recherche Scientifique (MFM), Région Aquitaine and Agence Nationale pour la Recherche (to DNA, MemoNeuro ANR2010-BLAN-1408-01). This work benefited from the support of the Biochemistry and Biophysics Facility of the Bordeaux Neurocampus funded by the LabEX BRAIN ANR-10-LABX-43 and the Animal Housing facility funded by Inserm and LabEX BRAIN ANR-10-LABX-43. The confocal analysis was done in the Bordeaux Imaging Center (BIC), a service unit of the CNRS-INSERM and Bordeaux University, member of the national infrastructure France BioImaging supported by the French National Research Agency (ANR-10-INBS-04).

---

**Figure S1. Spatial memory abilities of aged rats in the water-maze.** Learning performances are expressed as the mean latency (A,C) and mean distance travelled (B,D) to find the submerged platform for the first (A,B) and second (C,D) cohort of senescent rats.

**Table S1. DGNs produced in adolescent rats or embryos are not activated by spatial learning in aged rats**.

## References

Abrous, D. N., Koehl, M., and Le Moal, M. (2005) Adult neurogenesis: from precursors to network and physiology. Physiol Rev 85, 523–569.

Abrous, D. N. and Wojtowicz, J. M. (2015) Interaction between Neurogenesis and Hippocampal Memory System: New Vistas. Cold Spring Harb. Perspect. Biol. 7.

Ahlenius, H., Visan, V., Kokaia, M., Lindvall, O., and Kokaia, Z. (2009) Neural stem and progenitor cells retain their potential for proliferation and differentiation into functional neurons despite lower number in aged brain. J Neurosci 29, 4408–4419.

Aimone, J. B., Li, Y., Lee, S. W., Clemenson, G. D., Deng, W., and Gage, F. H. (2014) Regulation and Function of Adult Neurogenesis: From Genes to Cognition. Physiol Rev. 94, 991–1026.

Akers, K. G., Martinez-Canabal, A., Restivo, L., Yiu, A. P., De, C. A., Hsiang, H. L., Wheeler, A. L., Guskjolen, A., Niibori, Y., Shoji, H., Ohira, K., Richards, B. A., Miyakawa, T., Josselyn, S. A., and Frankland, P. W. (2014) Hippocampal neurogenesis regulates forgetting during adulthood and infancy. Science 344 598–602.

Altman, J. (1962) Are new neurones formed in the brains of adult mammals? Science 135 1127–1128.

Alvarez, D. D., Giacomini, D., Yang, S. M., Trinchero, M. F., Temprana, S. G., Buttner, K. A., Beltramone, N., and Schinder, A. F. (2016) A disynaptic feedback network activated by experience promotes the integration of new granule cells. Science 354 459–465.

Bettio, L. E. B., Rajendran, L., and Gil-Mohapel, J. (2017) The effects of aging in the hippocampus and cognitive decline. Neurosci. Biobehav. Rev. 79, 66–86.

Clelland, C. D., Choi, M., Romberg, C., Clemenson, G. D., Jr., Fragniere, A., Tyers, P., Jessberger, S., Saksida, L. M., Barker, R. A., Gage, F. H., and Bussey, T. J. (2009) A functional role for adult hippocampal neurogenesis in spatial pattern separation. Science 325 210–213.

Desjardins, S., Mayo, W., Vallee, M., Hancock, D., Le, M. M., Simon, H., and Abrous, D. N. (1997) Effect of aging on the basal expression of c-Fos, c-Jun, and Egr-1 proteins in the hippocampus. Neurobiol. Aging 18, 37–44.

Diaz-Moreno, M., Armenteros, T., Gradari, S., Hortiguela, R., Garcia-Corzo, L., Fontan-Lozano, A., Trejo, J. L., and Mira, H. (2018) Noggin rescues age-related stem cell loss in the brain of senescent mice with neurodegenerative pathology. Proc. Natl. Acad. Sci. U. S. A 115 11625–11630.

Drapeau, E. and Abrous, D. N. (2008) Role of neurogenesis in age-related memory disorders. Aging Cell 7, 569–589.

Drapeau, E., Mayo, W., Aurousseau, C., Le Moal, M., Piazza, PV., and Abrous, D. N. (2003) Spatial memory performances of aged rats in the water maze predict levels of hipppocampal neurogenesis. Proc Natl Acad Sci 100 14385–14390.

Drapeau, E., Montaron, M. F., Aguerre, S., and Abrous, D. N. (2007) Learning-induced survival of new neurons depends on the cognitive status of aged rats. J Neurosci 27, 6037–6044.

Dupret, D., Fabre, A., Dobrössy, M., Panetier, A., Rodriguez JJ, Lemaire V, Oliet, S. H. R., Piazza, P. V., and Abrous, D. N. (2007) Spatial learning depends on both the addition and removal of new hippocampal neurons. PLOS Biology 5, 1683–1694.

Dupret, D., Revest, J. M., Koehl, M., Ichas, F., De, G. F., Costet, P., Abrous, D. N., and Piazza, P. V. (2008) Spatial relational memory requires hippocampal adult neurogenesis. PLoS ONE. 3, e1959.

Encinas, J. M. and Fitzsimons, C. P. (2017) Gene regulation in adult neural stem cells. Current challenges and possible applications. Adv. Drug Deliv. Rev. 120 118–132.

Eriksson, P. S., Perfilieva, E., Bjork-Eriksson, T., Alborn, A. M., Nordborg, C., Peterson, D. A., and Gage, F. H. (1998) Neurogenesis in the adult human hippocampus. Nat Med 4, 1313–1317.

Fan, X., Wheatley, E. G., and Villeda, S. A. (2017) Mechanisms of Hippocampal Aging and the Potential for Rejuvenation. Annu. Rev. Neurosci. 40, 251–272.

Fischer, W., Wictorin, K., Bjorklund, A., Williams, L. R., Varon, S., and Gage, F. H. (1987) Amelioration of cholinergic neuron atrophy and spatial memory impairment in aged rats by nerve growth factor. Nature 329 65–68.

Geinisman, Y., Toledo-Morrell, L., and Morrell, F. (1986) Loss of perforated synapses in the dentate gyrus: morphological substrate of memory deficit in aged rats. Proc Natl Acad Sci U. S. A 83, 3027–3031.

Gheidi, A., Azzopardi, E., Adams, A. A., and Marrone, D. F. (2013) Experience-dependent persistent expression of zif268 during rest is preserved in the aged dentate gyrus. BMC. Neurosci. 14, 100.

Glatigny, M., Moriceau, S., Rivagorda, M., Ramos-Brossier, M., Nascimbeni, A. C., Lante, F., Shanley, M. R., Boudarene, N., Rousseaud, A., Friedman, A. K., Settembre, C., Kuperwasser, N., Friedlander, G., Buisson, A., Morel, E., Codogno, P., and Oury, F. (2019) Autophagy Is Required for Memory Formation and Reverses Age-Related Memory Decline. Curr. Biol. 29, 435–448.

Gonzalez-Escamilla, G., Muthuraman, M., Chirumamilla, V. C., Vogt, J., and Groppa, S. (2018) Brain Networks Reorganization During Maturation and Healthy Aging-Emphases for Resilience. Front Psychiatry 9, 601.

Gray, D. T. and Barnes, C. A. (2015) Distinguishing adaptive plasticity from vulnerability in the aging hippocampus. Neuroscience 309 17–28.

Gross, C. G. (2000) Neurogenesis in the adult brain: death of a dogma. Nat Rev Neurosci 1, 67–73.

Gu, Y., Arruda-Carvalho, M., Wang, J., Janoschka, S. R., Josselyn, S. A., Frankland, P. W., and Ge, S. (2012) Optical controlling reveals time-dependent roles for adult-born dentate granule cells. Nat. Neurosci 15, 1700–1706.

Hernandez-Segura, A., Nehme, J., and Demaria, M. (2018) Hallmarks of Cellular Senescence. Trends Cell Biol. 28, 436–453.

Kempermann, G., Gast, D., Kronenberg, G., Yamaguchi, M., and Gage, F. H. (2003) Early determination and long-term persistence of adult-generated new neurons in the hippocampus of mice. Development 130 391–399.

Lemaire, V., Tronel, S., Montaron, M. F., Fabre, A., Dugast, E., and Abrous, D. N. (2012) Long-lasting plasticity of hippocampal adult-born neurons. J Neurosci 32, 3101–3108.

Mahmoudi, S., Xu, L., and Brunet, A. (2019) Turning back time with emerging rejuvenation strategies. Nat. Cell Biol. 21, 32–43.

Marrone, D. F., Ramirez-Amaya, V., and Barnes, C. A. (2011) Neurons generated in senescence maintain capacity for functional integration. Hippocampus.

Marrone, D. F., Ramirez-Amaya, V., and Barnes, C. A. (2012) Neurons generated in senescence maintain capacity for functional integration. Hippocampus 22, 1134–1142.

Martin-Suarez, S., Valero, J., Muro-Garcia, T., and Encinas, J. M. (2019) Phenotypical and functional heterogeneity of neural stem cells in the aged hippocampus. Aging Cell 18, e12958.

Mayo, W., Lemaire, V., Malaterre, J., Rodriguez, J. J., Cayre, M., Stewart, M. G., Kharouby, M., Rougon, G., Le Moal, M., Piazza, P. V., and Abrous, D. N. (2005) Pregnenolone sulfate enhances neurogenesis and PSA-NCAM in young and aged hippocampus. Neurobiol. Aging 26, 103–114.

McAvoy, K. M. and Sahay, A. (2017) Targeting Adult Neurogenesis to Optimize Hippocampal Circuits in Aging. Neurotherapeutics. 14, 630–645.

Micheli, L., D’Andrea, G., Ceccarelli, M., Ferri, A., Scardigli, R., and Tirone, F. (2019) p16Ink4a Prevents the Activation of Aged Quiescent Dentate Gyrus Stem Cells by Physical Exercise. Front Cell Neurosci. 13, 10.

Montaron, M. F., Drapeau, E., Dupret, D., Kitchener, P., Aurousseau, C., Le, M. M., Piazza, P. V., and Abrous, D. N. (2006) Lifelong corticosterone level determines age-related decline in neurogenesis and memory. Neurobiol. Aging 27, 645–654.

Montaron, M. F., Petry, K. G., Rodriguez, J. J., Marinelli, M., Aurousseau, C., Rougon, G., Le Moal, M., and Abrous, D. N. (1999) Adrenalectomy increases neurogenesis but not PSA-NCAM expression in aged dentate gyrus. Eur J Neurosci 11, 1479–1485.

Moreno-Jiménez, E., Flore-Garcia, M., Terreros-Roncal, J., Rabano, A., Cafini, F., Pallas-Bazarra, N., Avial, J., and Llorens-Martin, M. (2019) Adult hippocampal neurogenesis is abundant in neurologically healthy subjects and drops sharply in patients with Alzheimer’s disease. nature Medecine 25, 554–560.

Nyberg, L., Lovden, M., Riklund, K., Lindenberger, U., and Backman, L. (2012) Memory aging and brain maintenance. Trends Cogn Sci. 16, 292–305.

O’Keefe J, N. L. (1978) The hippocampus as a cognitive map. Oxford University Press: Oxford.

Ohline, S. M., Wake, K. L., Hawkridge, M. V., Dinnunhan, M. F., Hegemann, R. U., Wilson, A., Schoderboeck, L., Logan, B. J., Jungenitz, T., Schwarzacher, S. W., Hughes, S. M., and Abraham, W. C. (2018) Adult-born dentate granule cell excitability depends on the interaction of neuron age, ontogenetic age and experience. Brain Struct. Funct.

Penner, M. R., Roth, T. L., Barnes, C. A., and Sweatt, J. D. (2010) An epigenetic hypothesis of aging-related cognitive dysfunction. Front Aging Neurosci 2, 9.

Penner, M. R., Roth, T. L., Chawla, M. K., Hoang, L. T., Roth, E. D., Lubin, F. D., Sweatt, J. D., Worley, P. F., and Barnes, C. A. (2011) Age-related changes in Arc transcription and DNA methylation within the hippocampus. Neurobiol. Aging 32, 2198–2210.

Qiao, J., Zhao, J., Chang, S., Sun, Q., Liu, N., Dong, J., Chen, Y., Yang, D., Ye, D., Liu, X., Yu, Y., Chen, W., Zhu, S., Wang, G., Jia, W., Xi, J., and Kang, J. (2019) MicroRNA-153 improves the neurogenesis of neural stem cells and enhances the cognitive ability of aged mice through the notch signaling pathway. Cell Death. Differ.

Rapp, P. R. and Gallagher, M. (1996) Preserved neuron number in the hippocampus of aged rats with spatial learning deficits. Proc Natl Acad Sci USA 93, 9926–9930.

Sahay, A., Scobie, K. N., Hill, A. S., O’Carroll, C. M., Kheirbek, M. A., Burghardt, N. S., Fenton, A. A., Dranovsky, A., and Hen, R. (2011) Increasing adult hippocampal neurogenesis is sufficient to improve pattern separation. Nature 472 466–470.

Samson, R. D. and Barnes, C. A. (2013) Impact of aging brain circuits on cognition. Eur. J. Neurosci. 37, 1903–1915.

Schaie, K. W. and Willis, S. L. (2015) Handbook of the Psychology of Aging. Elsevier.

Schouten, M., Bielefeld, P., Garcia-Corzo, L., Passchier, E. M. J., Gradari, S., Jungenitz, T., Pons-Espinal, M., Gebara, E., Martin-Suarez, S., Lucassen, P. J., De Vries, H. E., Trejo, J. L., Schwarzacher, S. W., De Pietri, T. D., Toni, N., Mira, H., Encinas, J. M., and Fitzsimons, C. P. (2019) Circadian glucocorticoid oscillations preserve a population of adult hippocampal neural stem cells in the aging brain. Mol. Psychiatry.

Smith, L. K., He, Y., Park, J. S., Bieri, G., Snethlage, C. E., Lin, K., Gontier, G., Wabl, R., Plambeck, K. E., Udeochu, J., Wheatley, E. G., Bouchard, J., Eggel, A., Narasimha, R., Grant, J. L., Luo, J., Wyss-Coray, T., and Villeda, S. A. (2015) beta2-microglobulin is a systemic pro-aging factor that impairs cognitive function and neurogenesis. Nat. Med. 21, 932–937.

Smith, T. D., Adams, M. M., Gallagher, M., Morrison, J. H., and Rapp, P. R. (2000) Circuit-specific alterations in hippocampal synaptophysin immunoreactivity predict spatial learning impairment in aged rats. J Neurosci 20, 6587–6593.

Spalding, K. L., Bergmann, O., Alkass, K., Bernard, S., Salehpour, M., Huttner, H. B., Bostrom, E., Westerlund, I., Vial, C., Buchholz, B. A., Possnert, G., Mash, D. C., Druid, H., and Frisen, J. (2013) Dynamics of hippocampal neurogenesis in adult humans. Cell 153 1219–1227.

Trinchero, M. F., Buttner, K. A., Sulkes Cuevas, J. N., Temprana, S. G., Fontanet, P. A., Monzon-Salinas, M. C., Ledda, F., Paratcha, G., and Schinder, A. F. (2017) High Plasticity of New Granule Cells in the Aging Hippocampus. Cell Rep. 21, 1129–1139.

Trinchero, M. F., Herrero, M., Monzon-Salinas, M. C., and Schinder, A. F. (2019) Experience-Dependent Structural Plasticity of Adult-Born Neurons in the Aging Hippocampus. Front Neurosci. 13, 739.

Tronel, S., Belnoue, L., Grosgean, N., Revest, J. M., Piazza, P. V., Koehl, M., and Abrous, D. N. (2012) Adult-born neurons are necessary for extended contextual discrimination. Hippocampus 22, 292–298.

Tronel, S., Charrier, V., Sage, C., Maitre, M., Leste-Lasserre, T., and Abrous, D. N. (2015a) Adult-born dentate neurons are recruited in both spatial memory encoding and retrieval. Hippocampus.

Tronel, S., Lemaire, V., Charrier, V., Montaron, M. F., and Abrous, D. N. (2015b) Influence of ontogenetic age on the role of dentate granule neurons. Brain Struct. Funct. 220 645–661.

Trouche, S., Bontempi, B., Roullet, P., and Rampon, C. (2009) Recruitment of adult-generated neurons into functional hippocampal networks contributes to updating and strengthening of spatial memory. Proc. Natl. Acad. Sci. U. S. A 106 5919–5924.

Veyrac, A., Besnard, A., Caboche, J., Davis, S., and Laroche, S. (2014) The transcription factor Zif268/Egr1, brain plasticity, and memory. Prog. Mol. Biol. Transl. Sci. 122 89–129.

Villeda, S. A., Luo, J., Mosher, K. I., Zou, B., Britschgi, M., Bieri, G., Stan, T. M., Fainberg, N., Ding, Z., Eggel, A., Lucin, K. M., Czirr, E., Park, J. S., Couillard-Despres, S., Aigner, L., Li, G., Peskind, E. R., Kaye, J. A., Quinn, J. F., Galasko, D. R., Xie, X. S., Rando, T. A., and Wyss-Coray, T. (2011) The ageing systemic milieu negatively regulates neurogenesis and cognitive function. Nature 477 90–94.

