## Supplementary figures and images for "Resilience to cognitive aging is associated with responsiveness of dentate neurons generated throughout adult life"

### Supplemental Figure 1

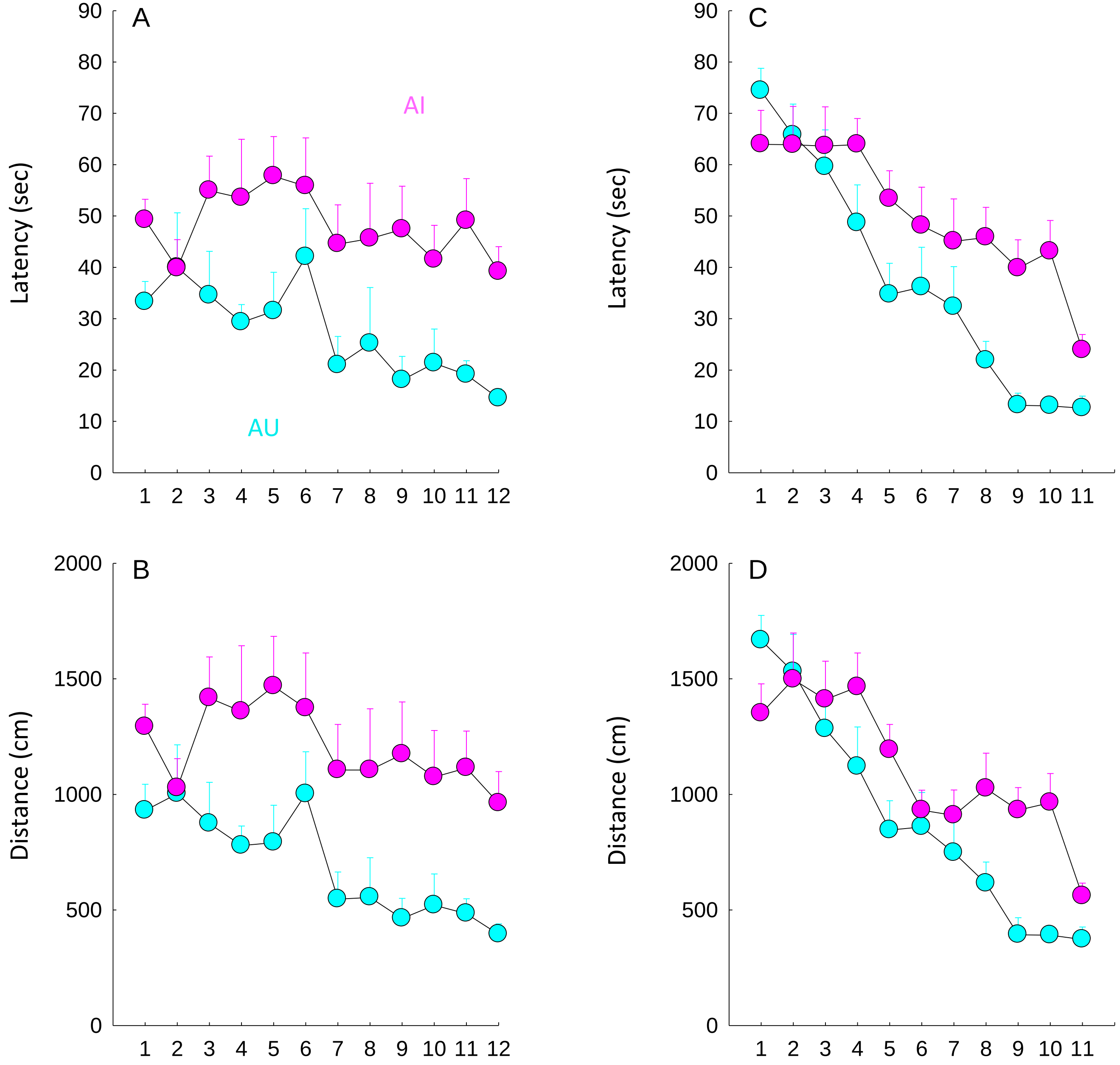
