## Supplemental Table 1 for "Resilience to cognitive aging is associated with responsiveness of dentate neurons generated throughout adult life"

| Experiments | Batch | Nb of Cldu-IR cells | Statistical analysis | % of CldU-Zif-268IR cells | Statistical analysis |
|---|---|---|---|---|---|
| **Ado**-DGNs | 3 | AU : 10496 ±1219<br>AI : 8626 ± 1027 | $t_8$=1.17, p=0.27 | AU : 2.08 ± 0.63<br>AI : 3.21 ± 0.76 | $t_8$=-1.14, p=0.29 |
| **Ado**-DGNs | 4 | AU : 6928 ± 2131<br>AI : 7724 ± 1473 | $t_8$=-0.31, p=0.77 | AU : 3.52 ± 1.51<br>AI : 2.62 ± 0.13 | $t_8$=0.59, p=0.57 |
| **Embryo**-DGNs | 5 | AU : 160510 ± 10375<br>AI : 149371 ± 25974 | $t_{11}$=0.42, p=0.68 | AU : 7.84 ± 0.63<br>AI : 6.74 ± 0.35 | $t_{11}$=1.45, p=0.17 |
